# Cytoskeletal assembly in axonal outgrowth and regeneration analyzed on the nanoscale

**DOI:** 10.1101/2021.11.04.467310

**Authors:** Max Hofmann, Lucas Biller, Uwe Michel, Mathias Bähr, Jan Christoph Koch

**Affiliations:** Department of Neurology, University Medical Center Göttingen, Göttingen, Germany

**Author notes:** Email addresses. Corresponding author: Jan C. Koch, MD.

**Keywords:** STED, cytoskeleton, regeneration, axon, membrane-associated periodic skeleton

## Abstract

The axonal cytoskeleton is organized in a highly periodic structure, the membrane-associated periodic skeleton (MPS), which is essential to maintain the structure and function of the axon. Here, we use stimulated emission depletion microscopy (STED) of primary rat cortical neurons in microfluidic chambers to analyze the temporal and spatial sequence of MPS formation at the distal end of growing axons and during regeneration after axotomy. We demonstrate that the MPS does not extend continuously into the growing axon but develops from patches of periodic β-spectrin II arrangements that grow and coalesce into a continuous scaffold. We estimate that the underlying sequence of nucleation, elongation, and subsequent coalescence of periodic β-spectrin II patches takes around 15 hours. Strikingly, we find that development of the MPS occurs faster in regenerating axons after axotomy and note marked differences in the morphology of the growth cone and adjacent axonal regions between regenerating and unlesioned axons. Moreover, we find that inhibition of the spectrin-cleaving enzyme calpain accelerates MPS formation in regenerating axons and increases the number of regenerating axons after axotomy. Taken together, we provide here a detailed nanoscale analysis of MPS development in growing axons.

## Introduction

The cytoskeleton plays a crucial role in the outgrowth and regeneration of axons. It enables the neuron to extend its axon to otherwise unknown lengths for cells. The cytoskeleton must withstand mechanical force and torsion but also needs to facilitate adaption and response to trauma or damage, like axonal lesions. Moreover, it builds the scaffold for the precise distribution of membrane channels and receptors along the axon and the basis for axonal transport, which are both essential for the function of the neuron.

Recent advances in super-resolution microscopy (SRM) have opened up a door for nanoscale observations of biological structures. In light of these advances, Xu et al. found that the cytoskeletal proteins actin, spectrin, and adducin form highly periodic structures in the axon, the membrane-associated periodic skeleton (MPS) (Xu, Zhong, & Zhuang, 2013). Originally discovered using stochastic optical reconstruction microscopy (STORM), this periodic arrangement was soon confirmed using other SRM-methods like stimulated emission depletion (STED) microscopy (Lukinavičius et al., 2014).

The MPS is prevalent in a broad range of neuronal cell types across different species (D’Este et al., 2016; He et al., 2016). It has also been shown in brain slices (Xu et al., 2013) and in living neurons (D’Este, Kamin, Göttfert, El-Hady, & Hell, 2015). Several studies have highlighted a broad range of involvements for the MPS in neuronal processes, like stabilization of axons (Hammarlund, Jorgensen, & Bastiani, 2007), electrical conductivity (Costa et al., 2020), lattice-like organization of proteins (Li et al., 2020; Vassilopoulos, Gibaud, Jimenez, Caillol, & Leterrier, 2019; Xu et al., 2013; Ruobo Zhou, Han, Xia, & Zhuang, 2019), and control of axonal diameter (Costa et al., 2020; Leite et al., 2016). It was established that the MPS consists of actin rings, made up of short actin filaments, capped by adducin, interconnected by α- and β-spectrin tetramers (Leite et al., 2016; Xu et al., 2013).

Several studies have suggested that the development of the MPS within the growing axon occurs in a proximal to distal pattern (Han, Zhou, Xia, & Zhuang, 2017; Lorenzo et al., 2019; Zhong et al., 2014). The MPS starts to form close to the soma at an early stage during axon development, at day in vitro (DIV) 2 in hippocampal rat neurons (Zhong et al., 2014). It then develops and progresses along the growing axon over time.

The role of the MPS in axonal degeneration has been studied previously (Unsain, Bordenave, et al., 2018; Wang et al., 2019). It was shown that axonal degeneration induced by trophic factor withdrawal led to a loss of the MPS (Unsain, Bordenave, et al., 2018; Wang et al., 2019), but this was independent of caspase apoptotic pathways (Wang et al., 2019). Interestingly, stabilization of F-actin using the drug cucurbitacin-B preserved the MPS and reduced axonal degeneration (Unsain, Bordenave, et al., 2018). In the last decade, mechanical axotomy through vacuum media aspiration in neurons cultured in microfluidic systems has proven to be a reliable model to study axonal degeneration and regeneration in vivo (Park, Vahidi, Taylor, Rhee, & Jeon, 2006; Taylor et al., 2005).

Despite this recent interest in the MPS and its development, the exact sequence and mechanism by which the MPS extends to newly formed parts of the outgrowing or regenerating axon remain unknown. Here, we provide a detailed nanoscale analysis of the MPS development in growing axons and the cytoskeletal changes upon regeneration after axotomy. We found that growing axons display a characteristic spatial and temporal increase of spectrin periodicity in a gradient from the distal growth cone towards the proximal axonal regions. We demonstrate that the building blocks in the formation of the MPS are periodic arrangements of β-spectrin II (periodic patches) that grow in size and number and eventually coalesce. In regenerating axons, the MPS develops faster and at more distal axon regions than physiologically outgrowing axons.

## Results

### Distribution of spectrin and tubulin along the axon and the growth cone in rat cortical neurons

To analyze the organization of the developing MPS in physiologically outgrowing (native) and in regenerating (after axotomy) axons, we cultured primary rat embryonic cortical neurons in compartmentalized microfluidic chambers (Taylor et al., 2005) (**Figure 1, A**). The neurons were seeded in the soma compartment and extended their axons through microgrooves (length 450 μm, width 3 μm) to the axonal compartment allowing the separate analysis of axons. Cells were fixed on day 11, immunostained against different cytoskeletal proteins, and analyzed using STED microscopy.

**Figure 1.**
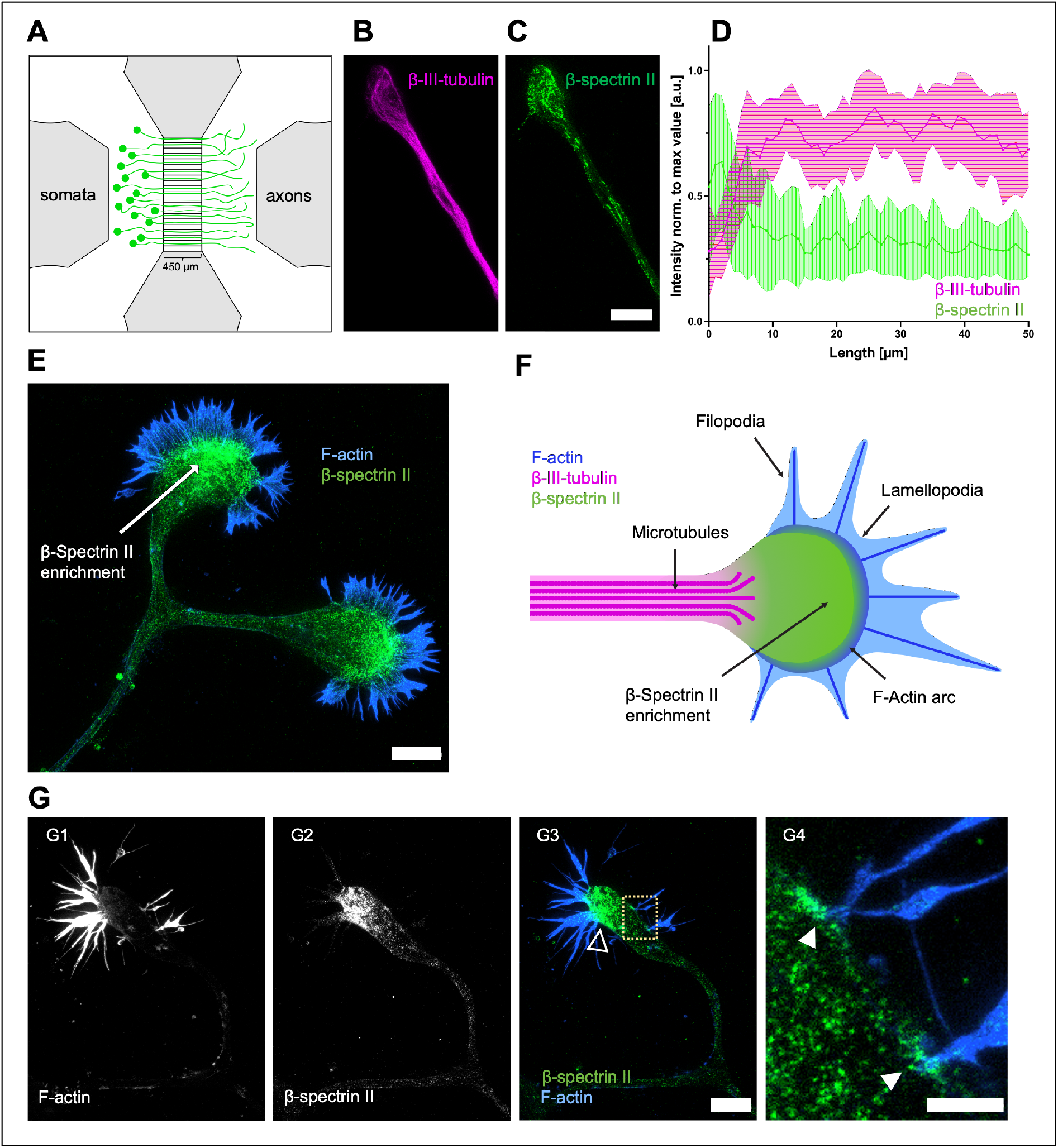
Distribution of β-spectrin II and β-III-tubulin along the axon and the growth cone in native rat cortical neurons. (**A**) Schematic drawing of a microfluidic chamber seeded with primary cortical neurons in the somatic department, transduced with AAV.EGFP virus. These neurons extend their axons through the microgrooves (length: 450 μm) into the axonal compartment within 11 days (green cells). (**B and C**) STED-images of cortical neurons stained for spectrin and tubulin. Note that spectrin is enriched in the growth cone while tubulin concentration is decreased. Scale bar 5 μm. (**D**) Quantification of line intensity scans along the axon, starting from the growth cone tip. Spectrin fluorescence intensity peaks at the growth cone, whereas the tubulin signal is lower compared to the axon. Data is represented as mean ± SD. At least 5 biological replicates were analyzed, 11 axons were analyzed in total. (**E**) Representative STED image of a growth cone stained for spectrin and F-actin. Spectrin is enriched close to the actin filaments. Scale bar 5 μm. (**F**) Schematic drawing of a growth cone, showing an enrichment of β-spectrin II in the apical central zone of the GC and microtubules splaying into the GC. (**G**) Representative STED image of a growth cone stained for F-actin and spectrin. Spectrin is enriched in the growth cone, in particular close to the base of filopodia (open arrowhead). **G4** shows an enlarged version of **G3**; closed arrowheads point to enriched spectrin at the base of filopodia. Scale bar 4 μm in **G1-3** and 1 μm in **G4**.

First, we studied the general spatial distribution of essential components of the axonal cytoskeleton along the native axon. We performed intensity line scans for β-spectrin II and β-III-tubulin along the growth cone (GC) and the adjacent axon (**Figure 1, B–D, Supplemental Figure 1**). The intensity of tubulin was highest along the distal axon shaft and significantly lower in the GC by 40% (0.44 ± 0.04 vs. 0.75 ± 0.02). In contrast, the intensity of spectrin was highest in the GC and was 65% lower along the axon (0.5 ± 0.04 vs. 0.3 ± 0.04). Both spectrin and actin intensity showed constant values further proximal up to 250 μm from the GC. We thus conclude that spectrin is strongly enriched in the GC and integrated into the axonal cytoskeleton in a lower concentration, while the opposite is true for tubulin.

The spectrin signal was strongest in the most apical central region and the transition region of the growth cone. In GCs co-stained for spectrin and F-actin, spectrin was enriched in the proximity to actin filaments. **Figure 1, E** and **F** show the location of the enrichment of spectrin in the GC relative to the F-actin-dominated filopodia. Especially at the base of F-actin bundles, the spectrin signal was enhanced and colocalized with actin. Exemplary micrographs are shown in **Figure 1, G**. This suggests a stabilizing or seeding function of spectrin for the outgrowing actin filaments in GCs.

### Development of the membrane-associated periodic skeleton in native rat cortical neurons

Zhong et al. reported that the MPS of hippocampal neurons starts to develop near the soma and matures along the axon towards the GC (Zhong et al., 2014). To better understand the process of MPS assembly in a growing axon, we asked how the MPS develops in the youngest part of the axon, i.e., the segment adjacent to GC. We, therefore, analyzed the first 250 μm of the axonal segment close to the GC (**Figure 2, A**). We divided the axon into 10 μm segments and evaluated for each respective segment the total length of the segment showing a periodic arrangement of spectrin with a distance of around 190 μm between spectrin signals. The respective length of axon showing MPS per respective segment was then divided by 10 μm. As depicted in **Figure 2, B**, there was a consistent, nearly 10-fold increase of axonal spectrin periodicity from the axonal GC base up to 250 μm towards the soma (first 10 μm: 7.5% ± 2.4%, 250 μm from GC: 68.5% ± 16.5%). A periodicity of 50% (5/10 μm of measured axon showed periodicity) was reached at a mean distance of 220 μm from the GC. With an average growth speed of 18.3 μm/hour in our culturing conditions, this distance corresponds to a maturation time of 12 hours for this axonal segment. A fully developed MPS (10/10 μm measured axon showed periodicity) was seen beyond distances of 250 – 350 μm from the GC. This distance corresponds to a maturation time of approximately 14 – 19 hours until an axonal segment contains a continuous MPS.

**Figure 2.**
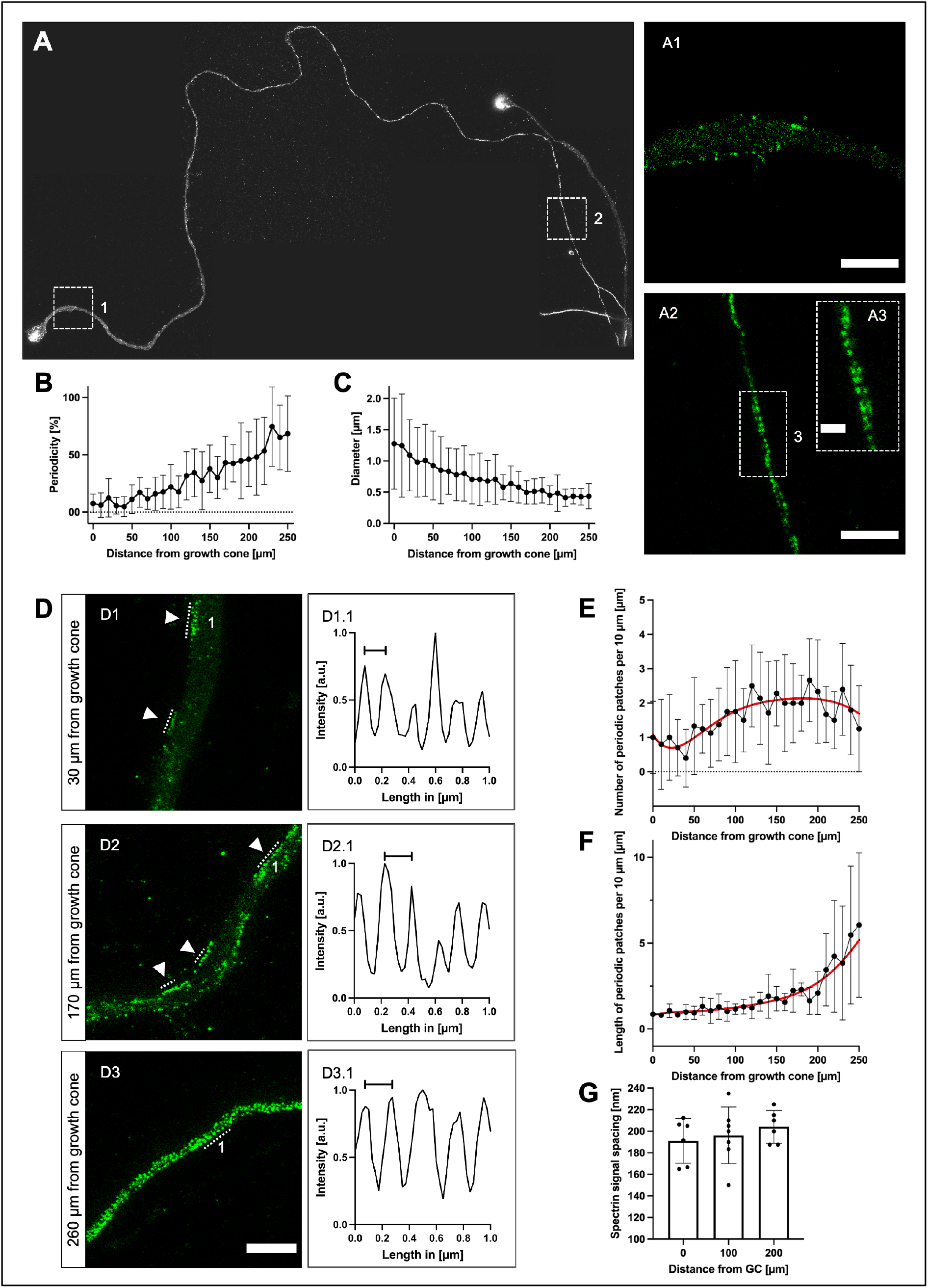
Development of the membrane-associated periodic skeleton along the growing axon. (**A**) Representative image of an axon with its growth cone, stained for spectrin, stitched from multiple STED images, up to 270 μm towards soma. **A1** shows an enlarged axon section at 10 μm distance from the growth cone, no periodicity is visible (scale bar 2 μm). **A2** shows an enlarged section of **A**, 270 μm from the growth cone (scale bar 2 μm). **A3** shows an enlarged section of **A2**, MPS is clearly visible (scale bar 0.5 μm). (**B**) Graphical display of the development of periodicity over the length of the axon, starting from the growth cone base. Periodicity is increasing linearly towards the soma. Data is represented as mean ± SD. At least 5 biological replicates were analyzed, 11 axons were analyzed in total. Because of limitations in following axons back for extended lengths, not all axons could be included at distal points. (**C**) Reduction of the axonal diameter over the length of the axon, starting from the growth cone base, contrariwise to (**B**). Data is represented as mean ± SD. At least 5 biological replicates were analyzed, 11 axons were analyzed in total. Because of limitations in following axons back for extended lengths, not all axons could be included at distal points. (**D**) Three sections of an axon stained for spectrin, showing different stages of MPS development, were imaged using STED microscopy. **D1** is 30 μm afar from the growth cone, **D2** 170 μm, and **D3** 260 μm. The arrowheads point to periodic patches, periodic arrangements of spectrin with at least three signals (scale bar 2 μm). **D1.1** shows the periodic plot of a line intensity scan along the numbered periodic patch in **D1**. The marked distance is 150 nm. **D2.1** shows a periodic line intensity scan along the numbered periodic patch in **D2**. The marked distance is 200 nm. **D3.1** shows the periodic plot of a line intensity scan along the numbered periodic patch in **D3**. The marked distance is 200 nm. (**E and F**) Quantification of parameters for the development of MPS along the axon, starting from the growth cone base. The number of periodic patches (**E**) increases, then decreases beyond 200 μm. The length of periodic patches (**F**) increases towards soma. Data is represented as mean ± SD. Red lines indicate estimated curve fits. At least 5 biological replicates were analyzed, 10 axons were analyzed in total. Because of limitations in following axons back for extended lengths, not all axons could be included at distal points. (**G**) Development of the spacing of spectrin tetramers in periodic patches in 10 μm long axonal segments at a distance of 0 μm, 100 μm, and 200 μm from the growth cone in native axons. Although a slight increase of spectrin tetramer spacing along the axon is visible, no significant changes were detected using ordinary one-way ANOVA, followed by Turkey’s multiple comparisons test. Bars represent mean ± SD, at least 5 biological replicates were analyzed, 11 axons were analyzed in total.

A role for the MPS in influencing the axonal diameter has been described in previous studies(Costa et al., 2020; Leite et al., 2016). We, therefore, analyzed the development of the axonal diameter along the axon. As **Figure 2, C** shows, there was a nearly three-fold decrease of axonal diameter along a distance of 250 μm from the GC (first 10 μm: 1.3 μm ± 0.2 μm, 250 μm from GC: 0.4 μm ± 0.1 μm). The finding that the youngest part of the axon shows the lowest periodicity, but greatest diameter, supports the idea of a progressive constriction of the axonal diameter by the MPS (Leite et al., 2016).

### The MPS develops in periodic patches

We noticed that spectrin periodicity does not extend continuously from proximal to distal but develops in the distal axon from multiple periodic spectrin patches extending and converging towards the soma (**Figure 2, D**). A “periodic spectrin patch” was defined as a continuous array of at least three spectrin-fluorescence signals inside the axon with a mean periodic distance of 150 – 250 nm. This distance was chosen based on the literature (Brown et al., 2015; Xu et al., 2013) and to take possible changes of spectrin tetramer spacing during maturation into account. The number and length of these periodic spectrin patches were measured in 10 μm segments along the growing axon starting at the base of the GC.

Short periodic spectrin patches appeared in the first 100 μm adjacent to the GC, with a density of 1 patch per 10 μm of axon (1.1 ± 0.1). The density increased to 2 patches per 10 μm between 100 – 200 μm from the GC (2.1 ± 0.1) and then slowly decreased again at distances >200 μm from the GC as single periodic spectrin patches extended and coalesced (1.8 ± 0.6 at 250 μm) (**Figure 2, E)**.

The mean length of the periodic spectrin patches in the first 100 μm adjacent to the GC was 0.6 μm (644 nm ± 96.4 nm), corresponding to 3 consecutive spectrin tetramers. Between 100 – 200 μm from the GC, the length of the periodic spectrin patches increased gradually (1525 nm ± 14.7 nm) and then exponentially beyond 200 μm to a mean length of 5.5 μm (5476 nm ± 179.6 nm) at 250 μm, corresponding to around 27 spectrin tetramers (**Figure 2, F**).

The average spacing of spectrin tetramers along the axon slowly increased from around 190 nm at the GC-base to 200 nm towards the soma (10 μm: 191.2 nm ± 8.5 nm, 100 μm: 196.2 nm ± 9.9 nm, and 200 μm from the GC: 204.2 nm ± 6.2 nm) (**Figure 2, G**).

Our data supports a model in which the MPS does not extend continuously into the distal outgrowing axon. Instead, the MPS assembles from multiple “MPS seeds” that grow and coalesce into a single scaffold. The formation of the MPS in an outgrowing axon can be divided into a nucleation phase with multilocular periodic spectrin patches developing along the axon, an elongation phase where single patches extend their size and the number of patches further increases, and a final coalescence phase where the MPS is closed (**Figure 3**).

**Figure 3.**
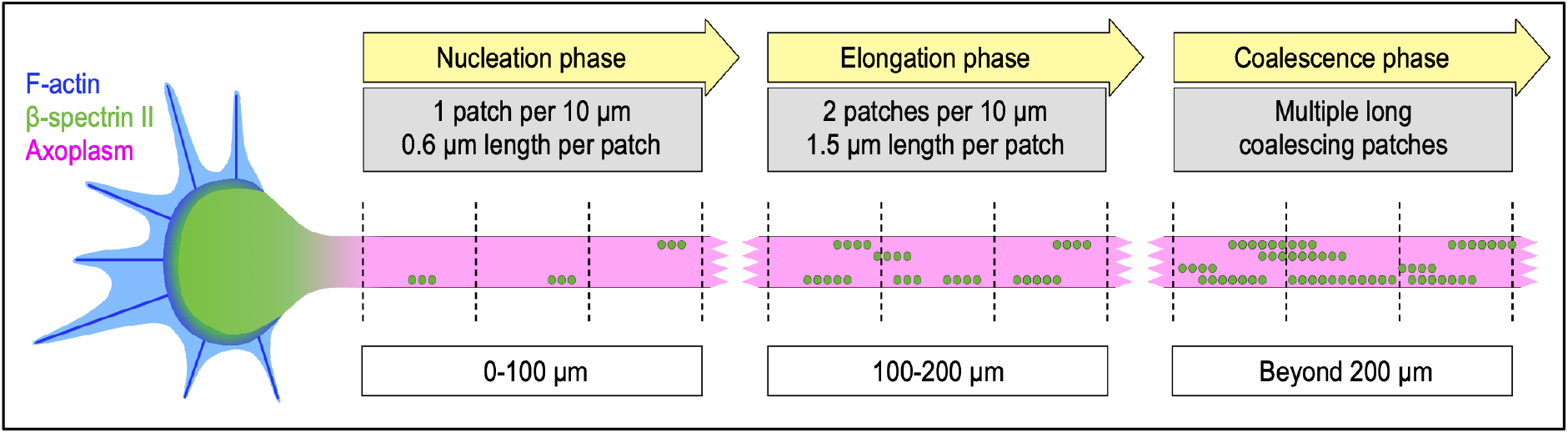
Phases of MPS development along the growing axon. Schematic drawing of the phases of MPS development along the growing axon, the growth cone is to the left. The dotted lines mark 10 μm intervals. β-Spectrin II molecules form periodic patches (symbolized by small green circles) close to the growth cone *(nucleation phase)*. These patches then increase in number and size *(elongation phase)*, coalesce during maturation, and form the final MPS *(coalescence phase)*.

### Spectrin periodicity is increased early after regeneration

Our next set of experiments aimed to analyze the morphological and cytoskeletal changes in regenerating axons following an axotomy. We performed axotomy by vacuum aspiration in microfluidic chambers and fixed the neurons and regenerating axons at different time points (2, 3, 4, 5, 6, and 24 hours) after axotomy. We then analyzed different morphological features of the GC and the axon and analyzed the development of the MPS in the axon. Because of the different lengths of the regenerating axons at different time points, we compared periodicity values only at the first 10 μm close to the GC at all time points. Each analyzed axon was traced over time to confirm its axotomy and to reliably exclude the few native axons growing through the microgrooves from the analysis (**Figure 4, A**; for further details, see methods section). Regeneration was first noticed at 2 hours after axotomy. Axons extended in length over the examined period.

**Figure 4.**
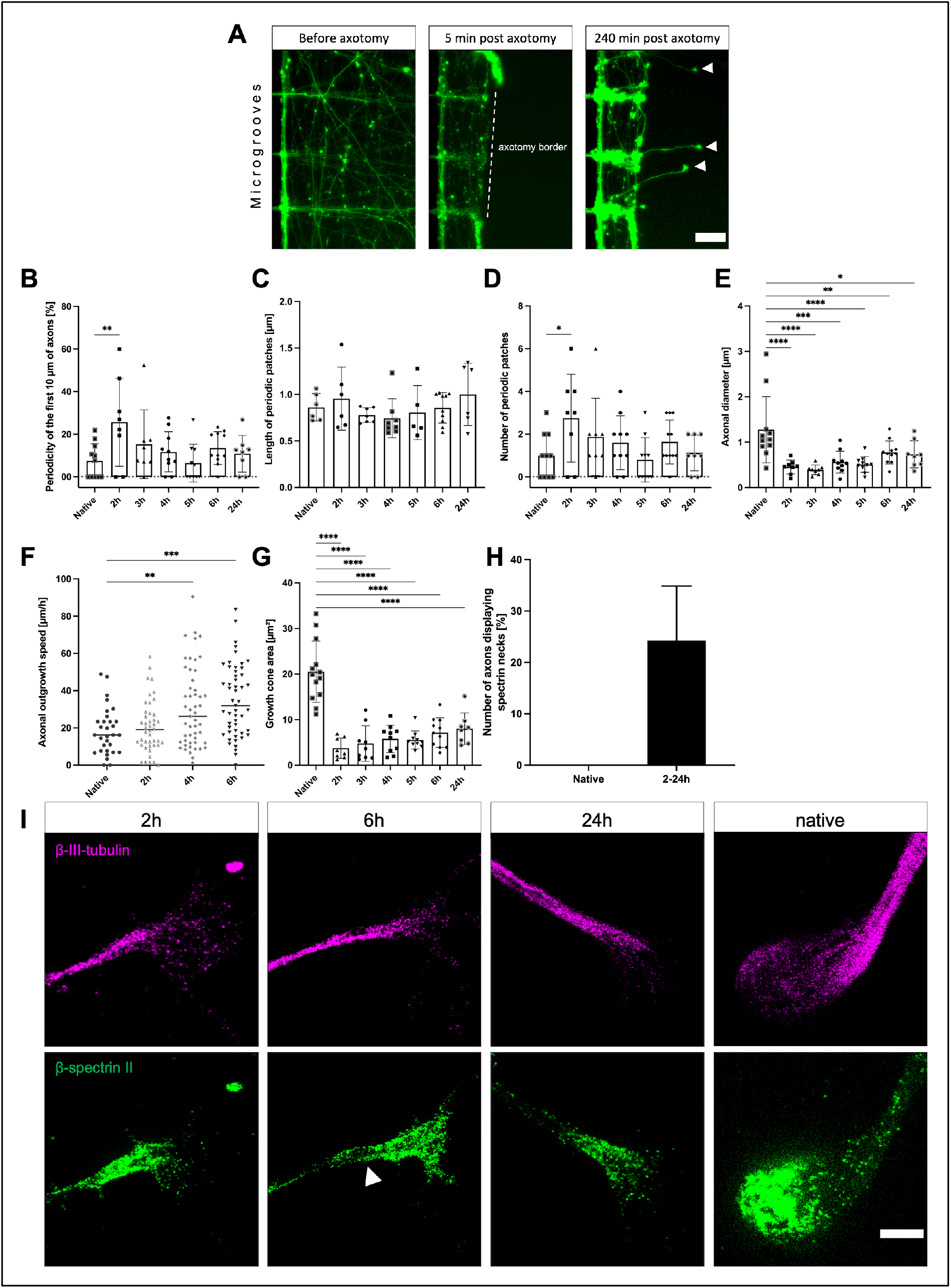
Comparison of growth cone morphology and β-spectrin II periodicity of the first 10 μm of axon proximal to the growth cone in native and regenerating neurons. (**A**) Graphical explanation of the method used for marking regenerating axons. Microscopic images of the same section of the axonal compartment were acquired before axotomy, five minutes after, and 2-24 hours after axotomy. For detailed information, see the methods section. White arrowheads indicate regenerating axons. Scale bar is 30 μm. **(B**) Comparison of periodicity in the first 10 μm of the axon, starting from the growth cone base. Data of different time points after axotomy and native axons are shown. The 2-hour time point shows a significant increase in periodicity. Bars represent mean ± SD. One-way ANOVA, followed by Dunnett’s multiple comparisons test, was performed. At least 8 axons were analyzed per condition, n=3, 67 axons were analyzed in total. (**C** and **D**) Comparison of the average length and number of periodic patches in the first 10 μm of axon adjacent to the growth cone. Data of different time points after axotomy and native axons are shown. No significant differences were detected for periodic patch length (**C**). The average number of periodic patches was significantly increased at the 2-hour time point, compared to native axons (**D**). One-way ANOVA was performed, followed by Dunnett’s multiple comparisons test. Bars represent mean ± SD. At least 8 axons were analyzed per condition, n=3, 67 axons were analyzed in total. (**E**) Comparison of axonal diameter in the first 10 μm of the axon, starting from the growth cone base. Data of different time points after axotomy and native axons are shown. Reduction of the axonal diameter of the different time points after axotomy is significant, compared to native axons, whereas only the 2-hour time point shows a significant increase in periodicity (**B**). Bars represent mean ± SD, one-way ANOVA, followed by Dunnett’s multiple comparisons test was performed. At least 8 axons were analyzed per condition, n=3, 67 axons were analyzed in total. (**F**) Comparison of axonal outgrowth speed in cortical neurons fixed at DIV 10. The 4-hour and 6-hour time points showed significantly increased outgrowth speed compared to native axons. The line represents mean, one-way ANOVA, followed by Dunnett’s multiple comparisons test was performed. At least 31 Axons were analyzed per condition, n=3, 177 axons were analyzed in total. (**G**), Comparison of growth cone area of different time points after axotomy and native axons, quantified in spectrin-stained axons. The growth cone area of regenerating axons is significantly reduced compared to native axons. Bars represent mean ± SD; one-way ANOVA, followed by Dunnett’s multiple comparisons test, was performed. At least 8 axons were analyzed per condition, n=3, 67 axons were analyzed in total. (**H**) Comparison of regenerating axons of combined time points and native axons displaying spectrin necks. Native axons did not display spectrin necks, whereas regenerating axons showed variating percentages of spectrin necks. A spectrin neck was defined as a visible increase of spectrin signal at the axonal segment directly connected to the growth cone, not longer than 3 μm. At least 8 axons were analyzed per condition, n=3, 67 axons were analyzed in total. (**I**) Representative STED images of growth cone sizes at different time points after axotomy and native. Notice the size difference of native and regenerating axons. White arrowhead points to spectrin neck. Scale bar is 2 μm.

Surprisingly, regenerating axons developed thiree MPS much faster than the native axons with a significant, almost 3.5-fold higher spectrin periodicity in the first 10 μm at 2 hours after axotomy in regenerating axons compared to native axons (**Figure 4, B**) (25.7% ± 7.3% vs. 7.5% ± 2.4%).

As **Figure 4, C shows**, the length of periodic patches was not significantly increased in regenerating axons, compared to native axons (0.96 μm ± 0.1 μm vs. 0.86 μm ± 0.06 μm). However, the number of periodic patches was increased almost three-fold at the 2-hour time point, compared to native axons (2.7 ± 0.7 vs. 1.0 ± 0.3; **Figure 4, D**).

We thus conclude that after axotomy, the axons transiently form more periodic spectrin patches in the outgrowing part of the axon resulting in a higher degree of periodicity and a quicker nucleation phase compared to native axons. Interestingly, at later time points after axotomy, the lesioned axons then resembled more and more the characteristics of the native axons with regards to their periodicity and number of periodic patches.

The axonal diameter of the segment adjacent to the GC was decreased in regenerating axons compared to native axons (**Figure 4, E**). At 2 hours after axotomy, the axonal diameter was reduced almost three-fold (2 hours: 0.5 μm ± 0.1 μm vs. native: 1.3 μm ± 0.2 μm). The axonal diameter increased in the following hours but was still significantly reduced almost two-fold up to 24 hours after the axotomy (24 hours: 0.7 μm ± 0.1 μm).

To test whether the outgrowth speed of regenerating axons was faster or not, we performed time-lapse microscopy and measured the outgrowth velocity of the GC in the time-lapse video. As **Figure 4, F** shows, we saw a significant increase of outgrowth velocity 4 and 6 hours after axotomy but not 2 hours after axotomy (native: 18.3 μm/h ± 2.3 μm/h, 2 hours: 20.5 μm/h ± 2.1 μm/h, 4 hours: 30.8 μm/h ± 3.0 μm/h, and 6 hours after axotomy: 34.5 μm/h ± 2.7 μm/h). Thus, the outgrowth velocity is not correlated to the higher spectrin periodicity.

To analyze whether the newly formed GC after axotomy differed morphologically from the native one, we measured the area of native and regenerating GCs of different time points. As **Figure 4, G** shows, the area of regenerating GCs at 2 hours after axotomy was reduced five-fold, compared to native axons. The GC area increased by two times over 24 hours but did not reach native size in this timespan (2 hours: 3.7 μm2 ± 0.8 μm2, 24 hours: 8.0 μm2 ± 1.2 μm2, and native axons: 20.5 μm2 ± 1.9 μm2). We observed a similar reduction regarding the length and width of the GCs; for detailed information, see **Supplemental Figure 2**.

Some axons showed an enrichment of spectrin close to the GC, which we will refer to as “spectrin neck”. A spectrin neck was defined as a visible increase of spectrin signal at the axonal segment directly connected to the GC, not longer than 3 micrometers. A fraction of 24,3% of regenerating axons showed spectrin necks compared to none of the native axons (**Figure 4, H** and **I**).

To sum up the results of the axotomy experiments, regenerating axons compared to native axons exhibit a transiently higher degree of spectrin periodicity based on an increased periodic spectrin patch density, a persistent smaller axon diameter, and a reduced growth cone size. Outgrowth velocity is increased at 4 and 6 hours after axotomy in the regenerating axons. A substantial number of regenerating axons show an accumulation of spectrin at the base of the GC.

### Inhibition of calpain increased periodicity and the number of regenerating axons shortly after axotomy

Since spectrin periodicity was increased early in regenerating axons, we wondered how a more stabilized spectrin cytoskeleton might influence the growth behavior of axons. To stabilize spectrin and putatively further increase periodicity after axotomy, we treated the neurons with the calpain inhibitor calpeptin (10 μM dissolved in dimethylsulfoxide (DMSO) or DMSO alone as control), performed an axotomy, and analyzed the axons 2 hours after axotomy (**Figure 5, A** and **B**). Calpain is a spectrin cleavage enzyme that is activated early after axotomy leading to a destruction of spectrin structures (Zhang et al., 2016).

**Figure 5.**
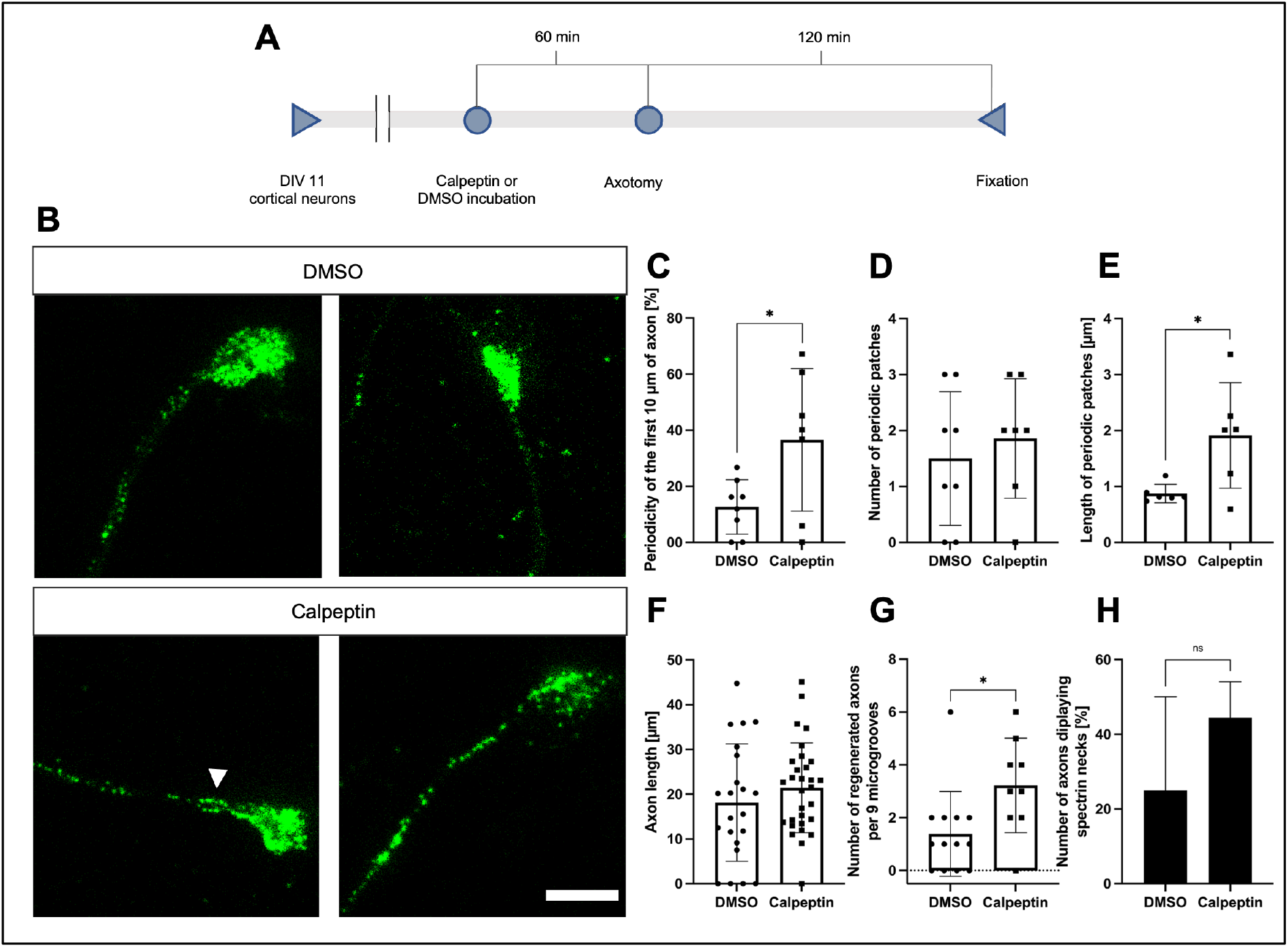
Effects of calpeptin treatment on MPS stability and regeneration of axons after axotomy. (**A**) Schematic display of used test setup. 10 μM of calpeptin or equal amounts of DMSO were used for incubation. STED imaging was performed following fixation. (**B**) Representative STED images of cortical neurons treated with calpeptin or DMSO before axotomy, fixation after 2 hours, scale bar 2 μm. The arrowhead points to a spectrin neck at the base of the growth cone. (**C-E**) Quantification of parameters for MPS development along the axon in regenerating axons treated with calpeptin, compared to regenerating DMSO controls. Analyzed were the first 10 μm of axon close to the growth cone. Periodicity (**C**) and size of periodic patches (**E**) were increased in calpeptin-treated axons. Bars represent mean ± SD. Unpaired t test (**C** and **D**) or Kolmogorov-Smirnov test (**E**) was performed. At least 7 axons were analyzed per condition, n=3, 15 axons were analyzed in total. (**F** and **G**) Graphical display of axon length of regenerating neurons 2 hours after axotomy. (**F**) shows no significant increase in axon length, while (**G**) shows a significant increase in regenerating axons per 9 microgrooves. Bars represent mean ± SD. Non-parametric Mann-Whitney U test was performed. At least 22 axons were analyzed per condition, n=3, 52 axons were analyzed in total. (**G**) at least 27 microgrooves were analyzed per condition, n=3, 198 microgrooves were analyzed in total. (**H**) Quantification of axons, treated with DMSO or calpeptin, displaying spectrin necks. Calpeptin-treated axons displayed a higher percentage of growth cones with spectrin necks. A spectrin neck was defined as a clearly visible increase of spectrin signal at the axonal segment directly connected to the growth cone, not longer than 3 μm. At least 7 axons were analyzed per condition, n=3, 15 axons were analyzed in total.

The calpeptin treatment indeed led to a more than 3-fold increase in spectrin periodicity at 2 hours after axotomy in the treated axons compared to control (36.6% ± 9.6% vs. 12.7% ± 3.4%) (**Figure 5, C)**. Interestingly, the axonal diameter did not differ between both groups (**Supplemental Figure 3**).

The number of periodic spectrin patches was not significantly changed in calpeptin-treated neurons compared to control (1.9 ± 0.4 vs. 1.5 ± 0.4) (**Figure 5, D**). The length of the periodic spectrin patches, however, was increased by 2.5-times in calpeptin treated neurons, compared to DMSO treated neurons (1.6 μm ± 0.4 μm vs. 0.7 μm ± 0.2 μm) (**Figure 5, E**). The number of periodic patches was decreased in both conditions compared to the previous experiment. This inconsistency could be due to toxic effects of the DMSO.

The length of the regenerating segment of the axons was not significantly different between calpeptin or DMSO treated neurons (21.4 μm ± 1.8 μm vs. 18.1 μm vs. 2.8 μm) (**Figure 5, F**). Interestingly, the number of regenerating axons per 9 microgrooves (9 microgrooves are in the viewfinder at 20x magnification) was significantly higher in analyzed images of calpeptin-treated chambers, compared to DMSO control group (3.2 ± 0.6 vs. 1.4 ± 0.4) (**Figure 5, G**). Also, axons treated with calpeptin showed spectrin necks more often than in the control group (44.4% vs. 25%), although the difference was not significant (**Figure 5, H**).

Inhibition of spectrin cleavage thus significantly enhances the elongation phase of MPS formation, leading to increased spectrin periodicity without influencing the nucleation phase. This does not increase outgrowth velocity but enables more axons to regenerate within the analyzed time period.

## Discussion

In this study, we analyzed the formation and development of the MPS in the outgrowing axon. We report that the developing MPS is organized in periodic patches that grow in size and number and coalesce over time during its maturation. We provide further evidence that the MPS develops in a proximal to distal pattern. We also show that regenerating axons after axotomy acquire periodicity faster over a transient timespan.

We found an enrichment of spectrin inside the axonal GC in line with previous studies (Galiano et al., 2012; Levine & Willard, 1981; Matsuoka, Li, & Bennett, 2000; Tian et al., 2012). This enrichment was present in native GCs and regenerated GCs at different time points after axotomy. Our data show that within the GC, spectrin is often localized in close proximity to the base of filopodia. Actin filaments could therefore be anchored or stabilized in the spectrin cytoskeleton of the GC. Elevated spectrin levels in the GC could also resemble a pool of spectrin tetramers that are later incorporated into the growing axon.

The MPS has been found along the axon, in dendrites, the AIS, and the neck of dendritic spines (Unsain, Stefani, & Cáceres, 2018). In outgrowing axons, the MPS has been suggested to develop in a proximal to distal pattern (Han et al., 2017; Lorenzo et al., 2019; Zhong et al., 2014). We can specify that growing axons display a characteristic spatial and temporal increase of spectrin periodicity in a gradient from the distal growth cone towards the proximal axonal regions. We show that the MPS does not extend in a continuous pattern along the axon. Instead, multiple periodic patches are nucleated along the growing axon. This nucleation happens close to the GC, at a frequency of about one patch per 10 μm. These periodic patches double in number over a timespan of about 11 hours, then increase in size and conflate as maturation continues to form the mature MPS finally. This maturation happens gradually as the axon grows, resulting in a gradient from immature MPS at the GC base towards mature MPS in the mid-axon.

We suggest the GC or its neck as a possible nucleation site for three reasons: First, the GC sits on top of the growing axon where spectrin nucleation naturally starts. Second, our data shows periodic patches in the axonal segment adjacent to the GC. Third, spectrin is highly enriched within the GC. It could therefore function as a pool for the implementation of spectrin into the growing axon. Interestingly, the number of periodic patches also increases at distances of 100 – 200 μm from the GC. This increase could be explained by nucleation molecules that stay on the way as the axon extends after integration into the axon at the GC. Zhong et al. demonstrated that formation of the MPS in dendrites is dependent on the local concentration of spectrin and the presence of ankyrin B (Zhong et al., 2014). Surprisingly, we did not detect periodic patches or periodicity inside the GC, despite higher concentrations of spectrin and ankyrin B (Galiano et al., 2012). This points to other nucleation factors that might not be present in the GC but in the axonal segment close to the GC. Our findings, therefore, shine a new light on the role of the GC and its neck in the formation of the axonal MPS.

A role of the MPS in maintaining axonal diameter has been described in previous studies (Costa et al., 2020; Leite et al., 2016). We show here that the axonal diameter decreases while the MPS is formed. The finding that the youngest part of the axon shows the lowest periodicity, but greatest diameter, supports the idea of a progressive constriction of the axonal diameter by the MPS (Leite et al., 2016). It has also been shown that the diameter of actin rings decreased over time in culture (Leite et al., 2016). Furthermore, knockout of α-adducin, also distributed periodically along the axon, increased the axonal diameter, without changes in the spacing of actin rings (Leite et al., 2016). Previous studies found recruitment of adducin into the MPS after DIV 8 (Zhong et al., 2014). It is also well known that non-muscle myosin II (NMII) has a role in regulating axonal diameter (Fan, Tofangchi, Kandel, Popescu, & Saif, 2017) and is distributed throughout the MPS (Costa et al., 2020; Vassilopoulos et al., 2019; R. Zhou et al., 2020). We found that stabilization of the MPS using calpeptin did not affect the axonal diameter in young, regenerating axons. This suggests that the role of both adducin and NMII in axonal diameter regulation is not directly linked to spectrin periodicity abundance and that spectrin plays an inferior role in diameter regulation. Interestingly, degeneration of the axon and its MPS through trophic factor withdrawal did also not lead to an increase in axonal diameter (Unsain, Bordenave, et al., 2018).

The role of the MPS in axonal degeneration has been studied previously (Unsain, Bordenave, et al., 2018; Wang et al., 2019). However, the MPS has not been studied in regenerating axons so far. Neuronal regeneration is an important process that needs to be activated in many neurological conditions in order to compensate for the disease-associated damage of nerve tissue, e.g., in spinal cord injury, stroke, or neurodegenerative diseases (Mahar & Cavalli, 2018).

We show here that axon morphology and the formation of the MPS in regenerating axons differ significantly from native axon outgrowth. Most strikingly, the axon diameter and growth cone size were significantly reduced in regenerating axons compared to native axons at all examined time points within 24h after axotomy. This reduction correlated with an increased outgrowth speed of the regenerating axons at 4 and 6 hours after axotomy.

At 2 hours after axotomy, spectrin periodicity and the number of spectrin patches were significantly increased in the newly formed axon but reached “normal” values at later time points. This suggests that the nucleation phase and most likely also the coalescence phase of MPS formation are enhanced early after axotomy. Since we accurately checked that all analyzed axons had been axotomized and evaluated only the newly formed part of the axons, we are confident that the observed alterations of spectrin MPS formation are characteristic of the regenerating axon and not confounded by residual proximal parts of the axotomized axon. A plausible explanation for the observations is that the molecular machinery necessary for nucleation and coalescence of spectrin patches is already in place in the more proximal part of the axon that remains after axotomy and can thus more quickly be employed for MPS formation. The decrease (i.e., normalization) of the number of periodic patches at later time points could be due to the increased outgrowth speed in the following hours, which would dilute a putatively still increased number of patches.

Also, spectrin itself is more abundant in the distal regenerating axon as compared to native axons. In around 25% of all regenerating axons, we saw an accumulation of spectrin adjacent to the GC, which we termed spectrin neck. Spectrin necks were never observed in axons without axotomy and thus specific for regenerating axons. The underlying mechanism for spectrin neck development might be increased local translation of spectrin RNA (Yoon, Zivraj, & Holt, 2009), altered axonal transport (Lorenzo et al., 2019), or local cleavage with increased spectrin reutilization (Wang et al., 2019).

Although hypothesized in a review by Leite and Sousa (2016), our finding of increased spectrin levels and periodicity in regenerating axons is particularly surprising, considering the degradation of the MPS in degenerating axons following axotomy or NGF-withdrawal, presented in previous studies (Unsain, Bordenave, et al., 2018; Wang et al., 2019). A possible explanation could be increased levels of free spectrin molecules after the axotomy. It has been shown that overexpression of spectrin increases periodicity in dendrites (Zhong et al., 2014). Since only a fraction of the axotomized axons regenerates while another fraction degenerates and only the regenerating axons were selected in our analysis, it can also be speculated that the observed features of the cytoskeleton are structural prerequisites that enable lesioned axons to regenerate after a lesion quickly.

It is well known that axotomy leads to a rapid calcium influx into the axoplasm and activation of the calcium-dependent protease calpain that induces the consecutive destruction of the axonal cytoskeleton (George, Glass, & Griffin, 1995; Knöferle et al., 2010). Inhibition of calpain, e.g., via calpeptin, has been shown to inhibit axonal degeneration after lesion, at least within the first hours after axotomy (George et al., 1995; Zhang et al., 2016). We thus analyzed the effects of calpeptin treatment on the spectrin cytoskeleton on the nanoscale. Calpeptin treatment indeed led to a significant increase of spectrin periodicity in regenerating axons. Interestingly, upon calpeptin treatment, the size of the periodic patches was increased, but not their number. Thus, calpain inhibition does not interfere with spectrin nucleation but enhances the elongation of periodic spectrin patches. The most likely explanation for this finding is a decreased degradation of spectrin molecules in the distal axon caused by calpain inhibition.

The increased and more rapidly evolving spectrin periodicity did not translate into an increased length of the regenerating axons. This is in line with previous studies that reported adverse effects of calpeptin on growth cone stability and axonal regeneration (Gitler & Spira, 1998). It is proposed that a constant turnover of the cytoskeletal molecules is essential for the highly dynamic growth cone and axonal elongation. However, we found an increased number of regenerating axons in the calpeptin-treated neurons. Thus, enhanced MPS-formation might set a basis for more axonal regeneration.

Taken together, we demonstrate here that the MPS is formed in the region 250 μm proximal from the GC in three phases: nucleation of periodic patches adjacent to the GC, subsequent elongation of periodic patches, and finally coalescence of periodic patches into a mature MPS. After a lesion, the outgrowth of regenerating axons is characterized by a greater speed, thinner axon diameter, smaller growth cones, and at least transiently an increased number of periodic patches and higher periodicity. Inhibition of the spectrin-severing protease calpain stabilized the MPS and enhanced the elongation phase. Future research should explore the underlying molecular mechanisms and test different substances that alter MPS dynamics.

## Materials and Methods

### Cell culture, viral transduction, and axotomy

The use of E18 Wistar rat embryos for isolation of cortical tissue was conducted according to the approved experimental animal licenses (33.9-42502-04-11/0408) issued by the responsible animal welfare authority (Niedersächsisches Landesamt für Verbraucherschutz und Lebensmittelsicherheit) and controlled by the local animal welfare committee of the University Medical Center Göttingen, Germany.

Primary cortical neurons were isolated from E18 Wistar rat embryos as described before (Sahu, Nikkilä, Lågas, Kolehmainen, & Castrén, 2019). In short, the dissected tissue was incubated with 1 ml trypsin (Sigma Aldrich, St. Louis, MO, USA; #T9935, 25.000 U/ml) for 15 min at 37 °C. Approximately 1 minute before the trypsinization was complete, 50 μl of DNase I (Roche, Basel, Swiss; #11284932001, 2000 U/mg) were added. The suspension was centrifuged shortly, and the supernatant was removed. Pellet was then resuspended in 1 ml fetal bovine serum (FBS; Biochrom, Berlin, Germany) and carefully triturated. If tissue clumps were still present, more DNase was supplemented. The supernatant was then transferred into a 15 ml falcon tube, and the remaining tissue was again triturated in 1 ml of media. The supernatant was transferred again, and the falcon tube was centrifuged. Neurons were resuspended in 500 μl Neurobasal medium (Gibco, Waltham, MA, USA), supplemented with 10% of FBS, 2% of 50x B27 Supplement (Gibco), 1% Penicillin-Streptomycin-Neomycin (Gibco), 0,5% holo-Transferrin human (Applichem, Darmstadt, Germany), and 0.25% L-glutamax (Gibco). Neurons were seeded at a density of 60,000 cells per chamber into the somatic compartment of a microfluidic chamber (MFC) (XONA microfluidics, Research Triangle Park, NC, USA). MFCs were mounted onto 35 mm FluoroDishes (World Precision Instruments, Inc, Sarasota, FL, USA), precoated with 0.1 mg/ml poly-L-ornithine (Sigma Aldrich, P3655) in borate buffer solution (Sigma Aldrich, P8638), and 0,1% laminin (Sigma Aldrich, L2020) as described previously (Zhang et al., 2016).

All cell cultures used for STED microscopy were transduced with an AAV 1/2, expressing EGFP (GenBank HQ416702.1), kindly gifted by Uwe Michel, Göttingen, as described before (Koch, Barski, Lingor, Bähr, & Michel, 2011). The stock solution had a titer of 5.6 x 107 transduction units (TU). Expression was driven by a human synapsin-1 promoter and a human h1 promoter. The virus was employed with a titer of 0.5 x 107 TU per chamber and added to the medium on DIV 4. Usually, a transduction rate of > 90% was achieved, with no significant toxicity. The medium was augmented from DIV 4 – 11 with 10 ng/μl of ciliary neurotrophic factor (CNTF, Pepro Tech, Rocky Hill, NJ, USA) and brain-derived neurotrophic factor (BDNF, Pepro-Tech). 10% of FBS was added to the medium in the first week but was omitted after DIV 6 to prevent the overpopulation of astrocytes. Cultured neurons were maintained with frequent medium exchange (every 2 to 3 days) and kept at 37 °C and 5% CO_2_. Cells looked healthy, and no signs of apoptosis were observed until DIV 11 following this protocol.

Axotomy was performed on DIV 11 by vacuum aspiration of the media in the axonal compartment as described elsewhere (Park et al., 2006). In short, the medium was aspirated using a cell culture pump (Schütt Labortechnik, Göttingen, Germany) until an air bubble passed through the axonal compartment, dissecting the axons. This procedure was repeated 2 – 4 times, and successful axotomy was assessed using a standard cell culture microscope (Zeiss, Jena, Germany). For treatment of neurons with calpeptin (Sigma Aldrich, #C8999), axons were incubated with 10 μM of calpeptin in DMSO (Applichem) or equal amounts of DMSO for 1 hour prior to the axotomy, administered only into the axonal compartment.

### Immunocytochemistry

All neuronal cultures were fixed on DIV 11. Axotomized neurons were fixed either 2, 3, 4, 5, 6, or 24 hours post-axotomy. Cells were washed with PBS (Applichem) and fixed with pure methanol (Applichem) for 10 minutes at −20°C. Afterward, fixed cells were washed three times with PBS and stored at 4°C in PBS if not immediately processed. Neuronal cultures used for staining actin were fixed for 1 hour using 4% paraformaldehyde (PFA; Applichem) in PHEM buffer, with 0.5% glutaraldehyde (Applichem), adapted from (Dent & Kalil, 2001).

For immunocytochemistry (ICC), MFCs were washed with PBS. If neurons were fixed with PFA, quenching with glycine (Applichem) and ammonium chloride (Applichem; both 100 mM in PBS) was performed to eliminate free aldehyde groups. After quenching (in case of PFA fixation) or initial washing (upon methanol fixation), cells were permeabilized with Triton X100 (0.3% in PBS; Applichem) for 5 min at room temperature (RT), MFCs were then incubated in blocking solution containing 2% bovine serum albumin (BSA) in PBS for 20 min at RT (Jackson Immuno Research, West Grove, PA, USA, #001-000-162). The blocking solution was removed, and primary antibodies were added for 1 hour at RT.

After the primary antibody reaction, suitable secondary antibodies or phalloidin STAR RED were added. Incubation time was set to 45 minutes and was performed at 37°C. If not immediately imaged, stained neurons were stored light-protected in PBS at 4°C.

Monoclonal primary antibodies directed against β-spectrin II were obtained from BD (BD Bioscience, Franklin Lakes, NJ, USA; mouse, #612563, 1:200), and rabbit β-III-tubulin antibodies were from Cell Signaling (Cambridge, UK; #5568S, 1:100). Polyclonal secondary antibodies goat anti-mouse Alexa594 were obtained from Invitrogen (Waltham, MA, USA; #A-11005, 1:100), and goat anti-rabbit STAR635p was from Abberior (#2-0012-007-2, 1:100, Göttingen, Germany) as was Phalloidin-STAR RED (#2-0205-011-7, 1:100). Calpeptin was obtained from Sigma (#C8999, 10 μM).

### Identification of regenerated axons

To identify and mark regenerated axons for STED microscopy, fluorescence live microscopy of neuronal cultures was performed. Cultures were imaged using an inverted fluorescence microscope (Axio Observer.Z1, Zeiss), equipped with an incubation chamber (Incubator PM 2000 RBT, PeCon, Erbach an der Donau, Germany), set to 37°C and 5% CO2. MFCs were placed inside the incubation chamber using a custom-made, 3D-printed template to seal the chamber against loss of CO2. Images were acquired at 20x magnification before axotomy, directly after, and at respective time points during regeneration (2, 3, 4, 5, 6, and 24 hours after axotomy). Using the “multiview” function of the Zen 2.5 software (blue edition, Zeiss), images were arranged side by side and aligned. After confirmation of a sufficient axotomy, the regenerating GCs that were not present before or shortly after the axotomy were marked and sorted out. Segments of the MFCs, showing neuronal somata on the axonal side were excluded. The microgrooves were numbered, and STED microscopy was performed only with axons previously identified as regenerating axons.

### Analysis of axonal outgrowth speed

Axonal outgrowth speed was assessed using time-lapse microscopy. In short, cells were cultured as described above. Several 10-minute-long time-lapse videos of the axonal compartment were captured before axotomy and 2, 4, and 6 hours after axotomy, at 40x magnification, using the setup described above. The position of the GC was marked in the first and last image of the video, and the distance traveled by the GC was then measured using the open-source freeware Fiji (Schindelin et al., 2009).

### Stimulated Emission Depletion (STED) microscopy

STED microscopy was performed to analyze MPS development in regenerating and native axons. At least two regenerated axons were recorded per condition and chamber, starting at the GC moving retrogradely towards the axotomy border. At 24 hours post-axotomy, axons were too long to be followed back to the axotomy border. Therefore, at least three images were acquired along the axon, starting from the GC. Native axons were imaged likewise. Two-color STED microscopy was performed using a quad scanning STED microscope (Abberior Instruments, Göttingen, Germany) equipped with an UPlanSApo 100x/1,40 Oil objective (Olympus, Tokyo, Japan). An IX83 Olympus microscope with 4-color LED illumination source and a monochrome widefield camera with a ½” CCD chip and 1280×960 pixels were used as a platform. Pixel size was set to 20 – 25 nm with a pinhole size of 0.7 airy units. Dwell times were set to 10 μs. Lasers with wavelengths of 561 nm and 640 nm were used for excitation of Alexa 594 or STAR 635p / STAR RED, respectively. Depletion was performed using a 775 nm pulsed STED laser. Images were acquired using the Imspector software (Abberior, Göttingen, Germany) (Schönle, 2006).

### Analysis of STED images

The brightness and contrast of the obtained STED images were adjusted using a custom-made macro and Fiji, mimicking the “auto-function” of the brightness and contrast tool, and converted into an 8-bit tagged image file format. If necessary, images of axon segments were stitched manually with MosaicJ (Thévenaz & Unser, 2007) upon deactivated smart color conversion, rotation, and blending. Images of the different cytoskeletal components were analyzed independently using Fiji. As for the evaluation of GC morphology (length, width, and area), the margin was outlined by applying the “polygon selection” tool. The length was determined by drawing a straight line from the GC base to the farthest point of the previously constructed polygon. The width was set to the longest straight within the polygon, being orthogonal to the length-line. A spline-fitted segmented line, with a thickness covering the axons width at the narrowest part, was drawn from the tip of the GC along the axon towards the axotomy border to plot intensity curves. To reduce data size and avoid outliers, we summarized the measurements for every 1 μm and normalized each dataset to its maximum. The axon itself was further evaluated regarding diameter and β-spectrin II periodicity in 10 μm segments, using the self-programmed “Segmenter” macro for the segmentation of nonlinear structures. The diameter was measured along the axon, starting at the GC neck, for every 10 μm. We calculated β-spectrin II periodicity by measuring every periodic patch in a segment and dividing the total length by 10μm. The number of periodic patches per segment and the average patch size were also assessed. One periodic patch was defined as at least three individual β-spectrin II-signals inside the axon with a mean distance of 150 – 250 nm between spectrin signals. This was assessed by performing intensity line scans along periodic patches, confirming the periodic distance of intensity peaks. Primary axons were analyzed likewise, but due to the absence of an axotomy border, the first 250 μm of the axon, starting at the GC base, were considered. To analyze the average spacing of spectrin tetramers, line intensity scans were performed in spectrin stains along periodic patches at distances of 0, 100, and 200 μm from the GC in all native axons. The distances between periodic signal peaks of spectrin were then measured. In co-stainings of actin and spectrin (**Figure 1, e** and **g**), actin is pseudo-colored using the CET-L15 colormap by Peter Kovesi (Kovesi, 2015) (colorcet.com).

### Statistics

Statistical evaluation was performed using GraphPad PRISM 9.1 (GraphPad Software, Inc, San Diego, CA, USA). Two groups were compared using unpaired Student’s t test (two-tailed). Non-parametric Mann-Whitney U or Kolmogorov-Smirnov tests were used if the assumptions for the t test were not met. Ordinary one-way ANOVA was conducted to compare two or more groups, followed by Dunnett’s multiple comparisons test. In general, mean values ± SEM are described unless otherwise noted, and at least three independent experiments were analyzed. Significant differences between compared groups are exhibited as follows: * p < 0.05, ** p < 0.01, *** p < 0.001, **** p < 0.0001.

## Acknowledgments

We thank Elisabeth Barski for excellent technical support and Alexander Böcker for numerous helpful comments on the manuscript. We also thank Stefan Hell and Stefan Stoldt for the provision of the STED facility and their continuous technical support with STED microscopy.

## Supplemental Figures

**Supplemental Figure 1.**
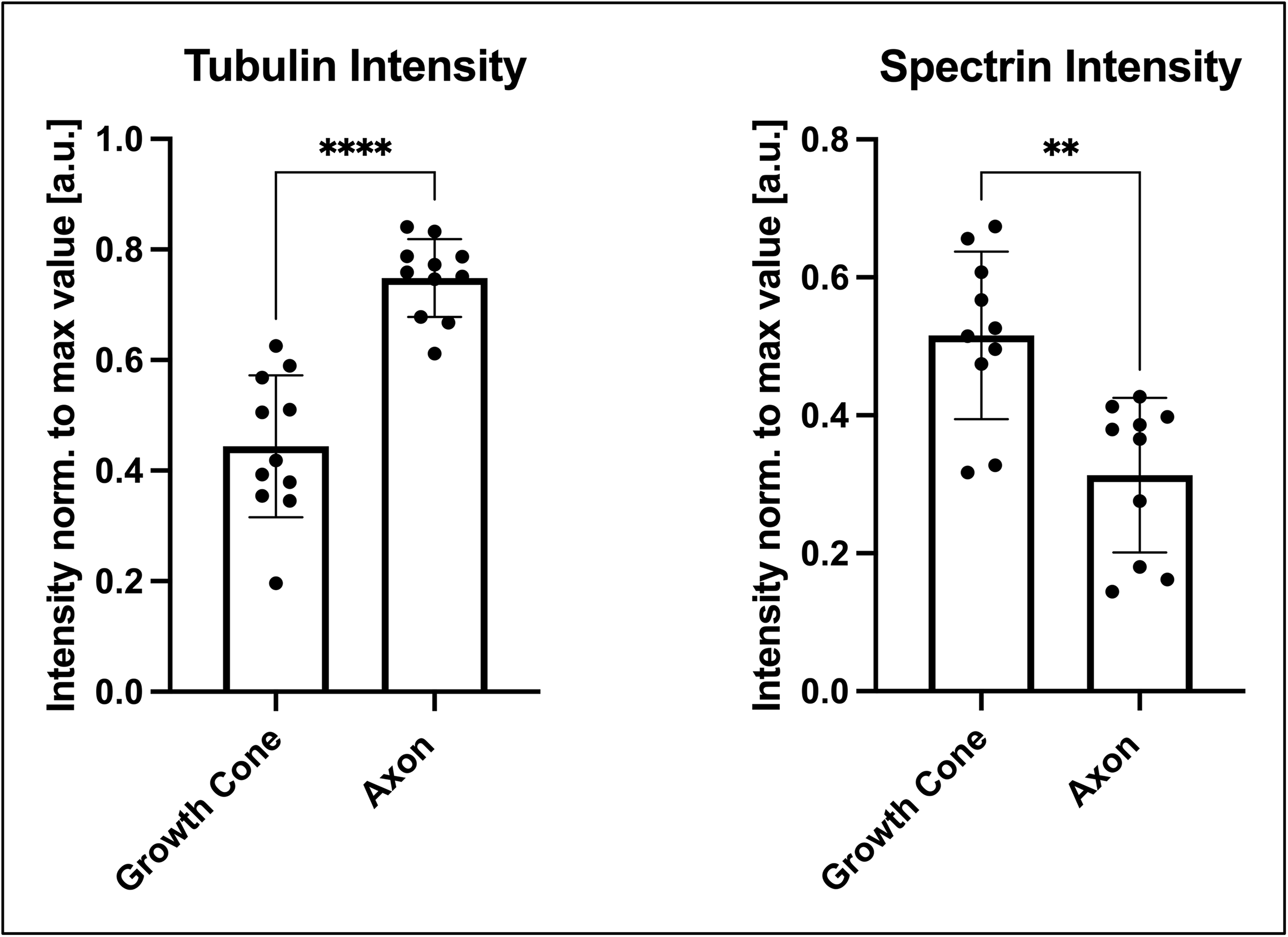
Distribution of β-spectrin II and β-III-tubulin along the axon and the growth cone in native rat cortical neurons. Quantification of 50 μm long line intensity scans along the axon, starting from the growth cone base. Due to the mean growth cone length, the first 7 μm of the line intensities represent the GCs, whereas the other 7 – 50 μm represent the axons. β-Spectrin II-fluorescence intensity peaks at the growth cone, whereas the β-III-tubulin signal is lower compared to the axon. Data is represented as mean ± SD. At least 4 biological replicates were analyzed, 11 axons were analyzed in total.

**Supplemental Figure 2.**
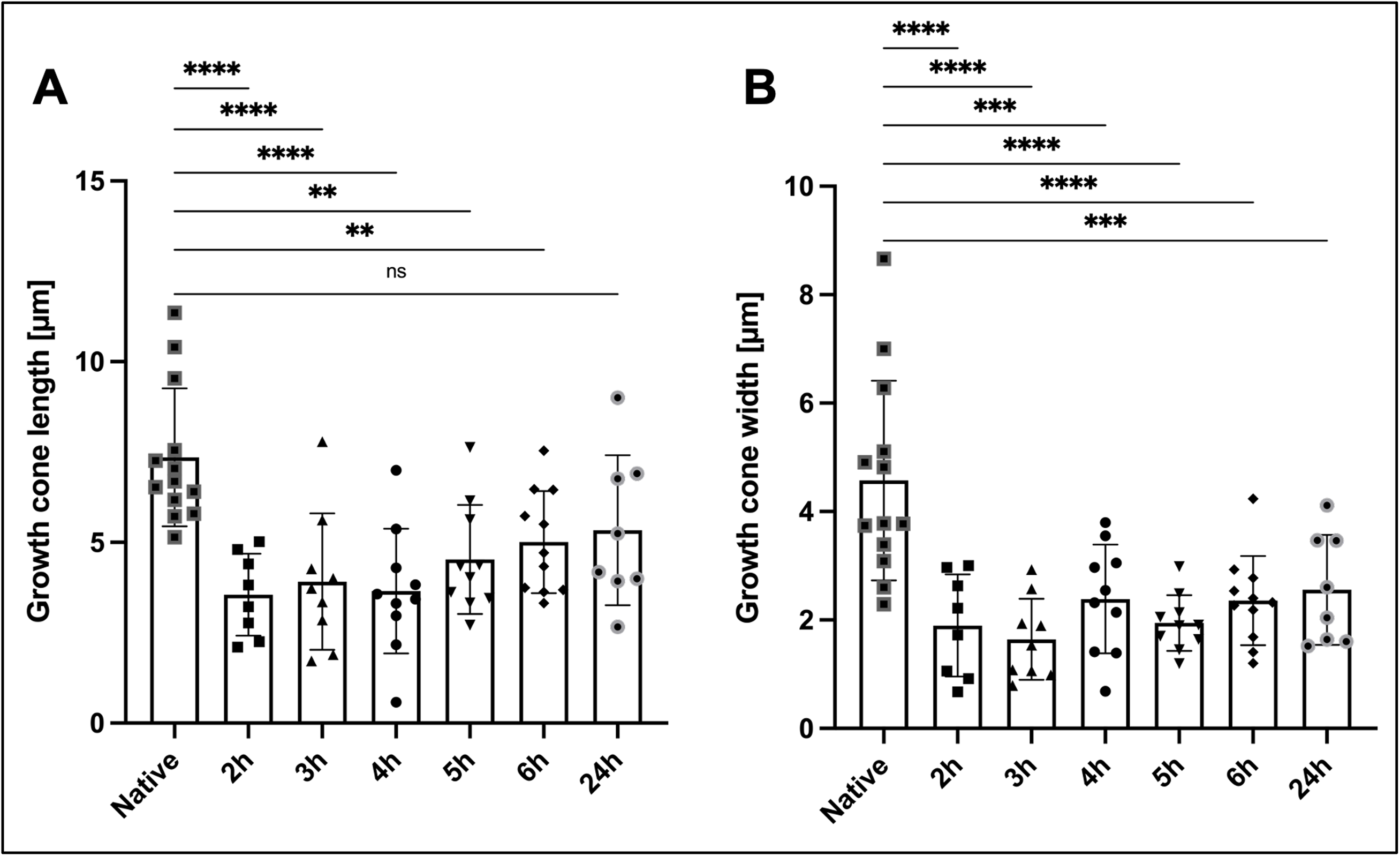
Difference of GC length and width of native and regenerating axons. Comparison of growth cone length (**A**), and growth cone width (**B**) of different time points after axotomy and native axons, quantified in spectrin-stained axons. The growth cone length and width of regenerating axons are significantly reduced compared to native axons, except for the growth cone width of the 24-hour time point. Bars represent mean ± SD; one-way ANOVA, followed by Dunnett’s multiple comparisons test, was performed. At least 8 axons were analyzed per condition, n=3, 67 axons were analyzed in total. (**A**) The length of regenerating GCs at 2 hours after axotomy was decreased two-fold compared to native axons. GC length was still significantly shorter up to 6 hours, but not 24 hours after axotomy (2 hours: 3.6 μm ± 0.4 μm, 6 hours: 5.0 μm ± 0.4 μm, 24 hours: 5.3 μm ± 0.7 μm, and native axons: 7.4 μm ± 0.5 μm). (**B**) The width of regenerating GCs was reduced by almost 2.5-times at 2 hours after axotomy, compared to native axons. This trend did not normalize over 24 hours, where GC width was still decreased by 1.8-times compared to native group (2 hours: 1.9 μm ± 0.3 μm, 24 hours: 2.6 μm ± 0.4 μm, and native axons: 4.6 μm ± 0.5 μm).

**Supplemental Figure 3.**
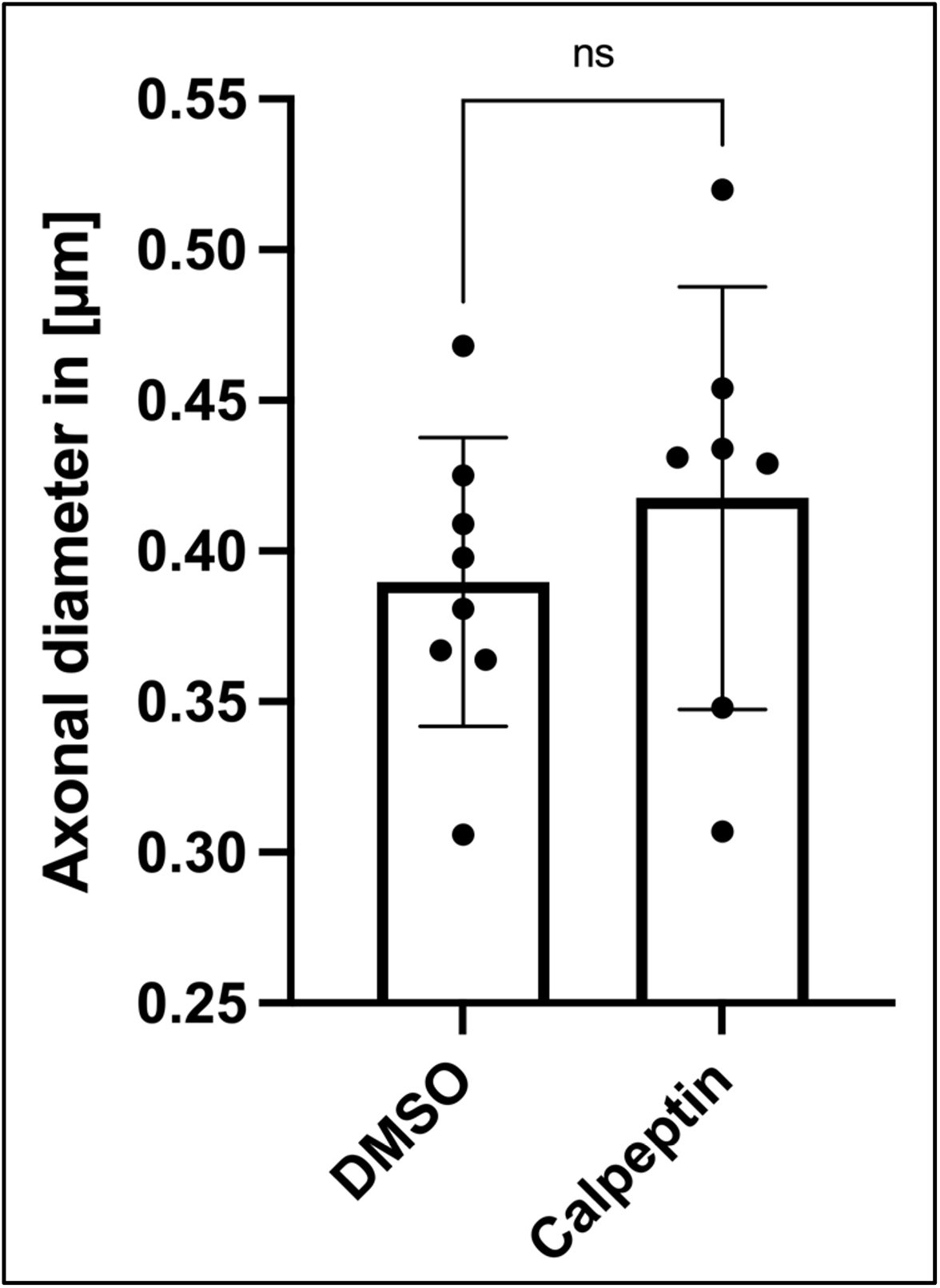
Diameter of the first 10 μm of axon adjacent to the growth cone in calpeptin or DMSO treated axons. Cells were cultured for 11 days. On DIV 11, calpeptin or DMSO was administered into the axonal compartment for 1 hour and axotomized afterward. Regenerating axons were fixed 120 minutes later and stained for spectrin and tubulin. Axonal diameter was then measured at 10 μm from the GC. Data is given as mean ± SD. At least 7 Axons were analyzed per condition, n=3, 15 axons were analyzed in total. No significant difference was detected according to unpaired t test.

## References

Brown, J. W., Bullitt, E., Sriswasdi, S., Harper, S., Speicher, D. W., & McKnight, C. J. (2015). The Physiological Molecular Shape of Spectrin: A Compact Supercoil Resembling a Chinese Finger Trap. PLoS Computational Biology, 11(6), e1004302. https://doi.org/10.1371/journal.pcbi.1004302

Costa, A. R., Sousa, S. C., Pinto-Costa, R., Mateus, J. C., Lopes, C. D. F., Costa, A. C.,…Sousa, M. M. (2020). The membrane periodic skeleton is an actomyosin network that regulates axonal diameter and conduction. ELife, 9, 1–20. https://doi.org/10.7554/eLife.55471

D’Este, E., Kamin, D., Göttfert, F., El-Hady, A., & Hell, S. W. (2015). STED Nanoscopy Reveals the Ubiquity of Subcortical Cytoskeleton Periodicity in Living Neurons. Cell Reports, 10(8), 1246–1251. https://doi.org/10.1016/j.celrep.2015.02.007

D’Este, E., Kamin, D., Velte, C., Göttfert, F., Simons, M., & Hell, S. W. (2016). Subcortical cytoskeleton periodicity throughout the nervous system. Scientific Reports, 6, 1–8. https://doi.org/10.1038/srep22741

Dent, E. W., & Kalil, K. (2001). Axon branching requires interactions between dynamic microtubules and actin filaments. Journal of Neuroscience, 21(24), 9757–9769. https://doi.org/10.1523/jneurosci.21-24-09757.2001

Fan, A., Tofangchi, A., Kandel, M., Popescu, G., & Saif, T. (2017). Coupled circumferential and axial tension driven by actin and myosin influences in vivo axon diameter. Scientific Reports, 7(1), 1–12. https://doi.org/10.1038/s41598-017-13830-1

Galiano, M. R., Jha, S., Ho, T. S. Y., Zhang, C., Ogawa, Y., Chang, K. J.,…Rasband, M. N. (2012). A distal axonal cytoskeleton forms an intra-axonal boundary that controls axon initial segment assembly. Cell, 149(5), 1125–1139. https://doi.org/10.1016/j.cell.2012.03.039

George, E. B., Glass, J. D., & Griffin, J. W. (1995). Axotomy-induced axonal degeneration is mediated by calcium influx through ion-specific channels. The Journal of Neuroscience?: The Official Journal of the Society for Neuroscience, 15(10), 6445–6452. https://doi.org/10.1523/JNEUROSCI.15-10-06445.1995

Gitler, D., & Spira, M. E. (1998). Real time imaging of calcium-induced localized proteolytic activity after axotomy and its relation to growth cone formation. Neuron, 20(6), 1123–1135. https://doi.org/10.1016/s0896-6273(00)80494-8

Hammarlund, M., Jorgensen, E. M., & Bastiani, M. J. (2007). Axons break in animals lacking β-spectrin. Journal of Cell Biology, 176(3), 269–275. https://doi.org/10.1083/jcb.200611117

Han, B., Zhou, R., Xia, C., & Zhuang, X. (2017). Structural organization of the actin-spectrin–based membrane skeleton in dendrites and soma of neurons. Proceedings of the National Academy of Sciences of the United States of America, 114(32), E6678–E6685. https://doi.org/10.1073/pnas.1705043114

He, J., Zhou, R., Wu, Z., Carrasco, M. A., Kurshan, P. T., Farley, J. E.,…Zhuang, X. (2016). Prevalent presence of periodic actin-spectrin-based membrane skeleton in a broad range of neuronal cell types and animal species. Proceedings of the National Academy of Sciences of the United States of America, 113(21), 6029–6034. https://doi.org/10.1073/pnas.1605707113

Knöferle, J., Koch, J. C., Ostendorf, T., Michel, U., Planchamp, V., Vutova, P.,…Lingor, P. (2010). Mechanisms of acute axonal degeneration in the optic nerve in vivo. Proceedings of the National Academy of Sciences, 107(13), 6064 LP–6069. https://doi.org/10.1073/pnas.0909794107

Koch, J. C., Barski, E., Lingor, P., Bähr, M., & Michel, U. (2011). Plasmids containing NRSE/RE1 sites enhance neurite outgrowth of retinal ganglion cells via sequestration of REST independent of NRSE dsRNA expression. FEBS Journal, 278(18), 3472–3483. https://doi.org/10.1111/j.1742-4658.2011.08269.x

Kovesi, P. (2015). Good Colour Maps: How to Design Them, 1–42. Retrieved from http://arxiv.org/abs/1509.03700

Leite, S. C., Sampaio, P., Sousa, V. F., Nogueira-Rodrigues, J., Pinto-Costa, R., Peters, L. L.,…Sousa, M. M. (2016). The Actin-Binding Protein α-Adducin Is Required for Maintaining Axon Diameter. Cell Reports, 15(3), 490–498. https://doi.org/10.1016/j.celrep.2016.03.047

Leite, S. C., & Sousa, M. M. (2016). The neuronal and actin commitment: Why do neurons need rings? Cytoskeleton, 73(9), 424–434. https://doi.org/10.1002/cm.21273

Levine, J., & Willard, M. (1981). Fodrin: axonally transported polypeptides associated with the internal periphery of many cells. The Journal of Cell Biology, 90(3), 631–642. https://doi.org/10.1083/jcb.90.3.631

Li, H., Yang, J., Tian, C., Diao, M., Wang, Q., Zhao, S.,…Zhong, G. (2020). Organized cannabinoid receptor distribution in neurons revealed by super-resolution fluorescence imaging. Nature Communications, 11(1). https://doi.org/10.1038/s41467-020-19510-5

Lorenzo, D. N., Badea, A., Zhou, R., Mohler, P. J., Zhuang, X., & Bennett, V. (2019). βiI-spectrin promotes mouse brain connectivity through stabilizing axonal plasma membranes and enabling axonal organelle transport. Proceedings of the National Academy of Sciences of the United States of America, 116(31), 15686–15695. https://doi.org/10.1073/pnas.1820649116

Lukinavičius, G., Reymond, L., D’Este, E., Masharina, A., Göttfert, F., Ta, H.,…Johnsson, K. (2014). Fluorogenic probes for live-cell imaging of the cytoskeleton. Nature Methods, 11(7), 731–733. https://doi.org/10.1038/nmeth.2972

Mahar, M., & Cavalli, V. (2018). Intrinsic mechanisms of neuronal axon regeneration. Nature Reviews. Neuroscience, 19(6), 323–337. https://doi.org/10.1038/s41583-018-0001-8

Matsuoka, Y., Li, X., & Bennett, V. (2000). Adducin: Structure, function and regulation. Cellular and Molecular Life Sciences, 57(6), 884–895. https://doi.org/10.1007/PL00000731

Park, J. W., Vahidi, B., Taylor, A. M., Rhee, S. W., & Jeon, N. L. (2006). Microfluidic culture platform for neuroscience research. Nature Protocols. https://doi.org/10.1038/nprot.2006.316

Sahu, M. P., Nikkilä, O., Lågas, S., Kolehmainen, S., & Castrén, E. (2019). Culturing primary neurons from rat hippocampus and cortex. Neuronal Signaling, 3(2). https://doi.org/10.1042/NS20180207

Schindelin, J., Arganda-Carrera, I., Frise, E., Verena, K., Mark, L., Tobias, P.,…Albert, C. (2009). Fiji - an Open platform for biological image analysis. Nature Methods, 9(7). https://doi.org/10.1038/nmeth.2019.Fiji

Schönle, A. (2006). Imspector Image Acquisition & Analysis Software, v0.1. Retrieved from http://www.imspector.de

Taylor, A. M., Blurton-Jones, M., Rhee, S. W., Cribbs, D. H., Cotman, C. W., & Jeon, N. L. (2005). A microfluidic culture platform for CNS axonal injury, regeneration and transport. Nature Methods, 2(8), 599–605. https://doi.org/10.1038/nmeth777

Thévenaz, P., & Unser, M. (2007). User-friendly semiautomated assembly of accurate image mosaics in microscopy. Microscopy Research and Technique, 70(2), 135–146. https://doi.org/10.1002/jemt.20393

Tian, N., Leshchyns’ka, I., Welch, J. H., Diakowski, W., Yang, H., Schachner, M., & Sytnyk, V. (2012). Lipid raft-dependent endocytosis of close homolog of adhesion molecule L1 (CHL1) promotes neuritogenesis. Journal of Biological Chemistry, 287(53), 44447–44463. https://doi.org/10.1074/jbc.M112.394973

Unsain, N., Bordenave, M. D., Martinez, G. F., Jalil, S., Von Bilderling, C., Barabas, F. M.,…Cáceres, A. O. (2018). Remodeling of the Actin/Spectrin Membrane-associated Periodic Skeleton, Growth Cone Collapse and F-Actin Decrease during Axonal Degeneration. Scientific Reports, 8(1), 1–16. https://doi.org/10.1038/s41598-018-21232-0

Unsain, N., Stefani, F. D., & Cáceres, A. (2018). The actin/spectrin membrane-associated periodic skeleton in neurons. Frontiers in Synaptic Neuroscience, 10(MAY), 1–8. https://doi.org/10.3389/fnsyn.2018.00010

Vassilopoulos, S., Gibaud, S., Jimenez, A., Caillol, G., & Leterrier, C. (2019). Ultrastructure of the axonal periodic scaffold reveals a braid-like organization of actin rings. Nature Communications, 10(1), 1–13. https://doi.org/10.1038/s41467-019-13835-6

Wang, G., Simon, D. J., Wu, Z., Belsky, D. M., Heller, E., O’Rourke, M. K.,…Zhuang, X. (2019). Structural plasticity of actin-spectrin membrane skeleton and functional role of actin and spectrin in axon degeneration. ELife, 8, 1–22. https://doi.org/10.7554/eLife.38730

Xu, K., Zhong, G., & Zhuang, X. (2013). Actin, spectrin, and associated proteins form a periodic cytoskeletal structure in axons. Science, 339(6118), 452–456. https://doi.org/10.1126/science.1232251

Yoon, B. C., Zivraj, K. H., & Holt, C. E. (2009). Local translation and mRNA trafficking in axon pathfinding. Results and Problems in Cell Differentiation, 48, 269–288. https://doi.org/10.1007/400_2009_5

Zhang, J. N., Michel, U., Lenz, C., Friedel, C. C., Köster, S., D’Hedouville, Z.,…Koch, J. C. (2016). Calpain-mediated cleavage of collapsin response mediator protein-2 drives acute axonal degeneration. Scientific Reports, 6(October), 1–15. https://doi.org/10.1038/srep37050

Zhong, G., He, J., Zhou, R., Lorenzo, D., Babcock, H. P., Bennett, V., & Zhuang, X. (2014). Developmental mechanism of the periodic membrane skeleton in axons. ELife, 3, 1–21. https://doi.org/10.7554/eLife.04581

Zhou, R., Han, B., Nowak, R., Lu, Y., Heller, E., Xia, C.,…Zhuang, X. (2020). Proteomic and functional analyses of the periodic membrane skeleton in neurons. BioRxiv. https://doi.org/10.1101/2020.12.23.424206

Zhou, Ruobo, Han, B., Xia, C., & Zhuang, X. (2019). Membrane-associated periodic skeleton is a signaling platform for RTK transactivation in neurons. Science, 365(6456), 929–934. https://doi.org/10.1126/science.aaw5937

